# Estimating functional EEG sources using topographical templates

**DOI:** 10.1101/2022.07.20.500772

**Authors:** Marlene Poncet, Justin Ales

**Affiliations:** University of St Andrews, School of Psychology and Neuroscience, United Kingdom

## Abstract

Electroencephalography (EEG) is a common and inexpensive method to record neural activity in humans. However, it lacks spatial resolution making it difficult to determine which areas of the brain are responsible for the observed EEG response. Here we present a new easy-to-use method that relies on EEG topographical templates. Using MRI and fMRI scans of 50 participants, we simulated how the activity in each visual area appears on the scalp and averaged this signal to produce functionally defined EEG templates. Once created, these templates can be used to estimate how much each visual area contributes to the observed EEG activity. We tested this method on extensive simulations and on real data. The proposed procedure is as good as bespoke individual source localization methods and has several strengths. First, because it does not rely on individual brain scans, it is inexpensive and can be used on any EEG dataset, past or present. Second, the results are readily interpretable in terms of functional brain regions and can be compared across neuroimaging techniques. Finally, this method is easy to understand, simple to use, and expandable to other brain sources. We thus expect it to be of wide interest to EEG users.

## Introduction

Electroencephalography (EEG) is a powerful tool to measure and study neuronal activity in the human brain. One of its major advantages is its very high temporal resolution. However, because EEG is recorded on the scalp, it represents the combined activity of multiple brain areas, making it difficult to determine the intracranial sources of the scalp signal. Various methods have been proposed to localize the brain sources of EEG activity (He et al., 2018; Michel & He, 2019; Michel & Murray, 2012). These source localization methods have been improved over decades, particularly with the introduction of anatomical and physiological constraints on the cortical sources (Dale & Sereno, 1993; Pascual-Marqui et al., 1994). Today, the best results are found with source localization methods that use realistic head models. These are created specifically for each individual from their MRI scans. It improves the reliability and precision of EEG source localization compared to previously used spherical head models, head models derived from a template MRI (such as MNI or Talairach) or from an averaged MRI dataset (Akalin Acar & Makeig, 2013; Baillet, Riera, et al., 2001; Brodbeck et al., 2011; Fuchs et al., 2002, 2007; Guggisberg et al., 2011; G. Wang et al., 2011).

Given that brain anatomy is highly variable across individuals, source localization methods are traditionally performed on an individual-basis. This also means that the retrieved brain sources are not the same across individuals and cannot be averaged easily. Although such anatomical localization is important in some circumstances (e.g., for localizing epileptic foci), most studies are interested in the neural mechanisms involved in a particular cognitive process or behavior at the group-level. That is, they are not interested in the anatomical brain sources per se but in identifying the functional brain area(s) involved in the process that is investigated. In line with this, a different approach, fMRI-informed EEG source imaging, has been developed where Regions of Interests (ROIs) are mapped out from fMRI scans using retinotopic mapping (Engel et al., 1994; Himmelberg et al., 2021; Wandell & Winawer, 2011) and/or fMRI localizers (Huk et al., 2002; Kanwisher et al., 1997; Saxe et al., 2006). Sources are then defined as belonging to a given ROI and because they represent the same brain function, they can be averaged across participants with different brain anatomy (this is common in fMRI when averaging BOLD response of voxels within ROIs). Another advantage of this method is that the activity between functionally equivalent sources can be compared across experiments using the same (EEG) or different (MEG, fMRI) technique. Besides, considerations about which electrodes to pick or to pool when analyzing the data become unnecessary. This approach has been very successful, especially in the field of visual perception (Ales, Appelbaum, et al., 2013; Ales & Norcia, 2009; Appelbaum et al., 2008; Cottereau et al., 2014; Cottereau, McKee, et al., 2012; Lauritzen et al., 2010; Palomares et al., 2012; Verghese et al., 2012; J. Wang & Wade, 2011). However, fMRI-informed EEG source imaging, like other methods, relies on obtaining MRI and fMRI scans to create individual realistic head models. Even if the head model of a participant can be used for multiple EEG experiments, the procedure is still expensive in terms of money, time, and computational load.

Here we present a new source localization method that uses topographies of EEG activity derived for multiple functionally defined visual brain area to recover the intracranial sources responsible for a given EEG scalp activity. This method substantially simplifies and reduces the costs of EEG source localization. It is inspired by the use of cortical (Cabezas et al., 2011; Evans et al., 2012) and functional atlases (Engell & McCarthy, 2013; Huang et al., 2019; Rosenke et al., 2020; Weiner et al., 2018; Zhen et al., 2017) to localize ROIs in new individuals. The implicit assumption in such an approach, shared by most neuroimaging procedures, is that the average of multiple participants is a good representation of the population. Following this idea, it should be possible to generate a scalp response from the average of multiple individuals that would represent the activity of a specific brain area. As the number of participants grows, the expected scalp response will converge towards the population average and not be dependent on the scalp response for any specific individual or the specific set of participants included in the study. This means that this expected response, or template, can be applied to any dataset. By creating multiple templates representing the activation of different ROIs, we can then determine how much each template contributes to the scalp EEG response.

EEG recordings are typically analyzed by averaging the signal across multiple participants to improve the signal-to-noise ratio (SNR). However, many EEG source imaging methods work by estimating the sources of individual subject’s data and then averaging across participants. This can result in poor quality estimates because source localization methods are typically sensitive to SNR. By effectively trying to sharpen the blurred scalp data, these methods also amplify noise. Source localization using group-informed EEG source imaging is better compared to retrieving sources at the individual level (Lim et al., 2017). By averaging across participants before source localization, SNR will be higher and can result in better source localization accuracy. The method that we propose uses the average of expected scalp responses for a set of ROIs (EEG templates) and the average of the recorded EEG signal to determine the contribution of each ROI to this EEG signal.

In contrast to other source localization methods using individual anatomic and functional MRI scans, the template method is based on the EEG templates that are created a priori and do not depend on specific participant data. Although we use fMRI and MRI data to create the EEG templates, once created, there is no need for additional scans. This method thus eliminates scan-related costs, is also much faster (as there is no need for additional MRI or fMRI data processing) and has the advantage that it can be applied to any past and future EEG recordings.

We expect this method to be useful for a wide range of research groups. Our fitting procedure is implemented using regularized linear regression, a tool widely used and understood, making this procedure relatively straightforward to apply in novel contexts. Our overall approach is easily implemented using the set of functions and EEG templates that we have made available at https://github.com/aleslab/eegSourceTemplateMatching.

## Methods

The template method that we propose is based on using EEG topographies that represent the activity of functional brain areas. We first describe how we created these topographies and then how we use them to recover brain sources from EEG scalp activity. To assess the validity of this method, we compare it with fMRI-informed source localization methods using simulated and real data.

### Creation of functionally defined EEG templates

The templates were constructed using boundary element forward models which define how the activity of a neural source propagates to each of the EEG electrodes at the surface of the scalp. To account for the variability in brains across individuals, we used 50 participants’ pre-analyzed structural and functional MRI scans collated from several experiments. These were originally collected with ethical approval from UCSF, The Smith-Kettlewell Eye Research Institute, and Stanford University (Ales et al., 2010; Cottereau, Ales, et al., 2012; Cottereau, McKee, et al., 2012; Lim et al., 2017).

The data included the definition of a source space and the surface boundaries for skin, skull, cerebrospinal fluid for each participant, co-registered with the positions of 128 EEG electrodes. In addition, 18 visual ROIs (V1-L, V1-R, V2v-L, V2v-R, V2d-L, V2d-R, V3v-L, V3v-R, V3d-L, V3d-R, V4-L, V4-R, V3A-L, V3A-R, LOC-L, LOC-R, MT-L, MT-R; where L=left, R=right, d=dorsal, v=ventral,) were defined based on high-resolution T1 whole-head anatomical MRI scans combined with functional MRI scans for each participant. Details of the analyses can be found in previous studies (Cottereau, Ales, et al., 2012; Cottereau, McKee, et al., 2012; Lim et al., 2017). In brief, the brain grey matter was delineated using FreeSurfer from a structural scan of each participant. The tessellation of the grey matter defined the source space and consisted in 20,484 regularly spaced vertices. The inner skull, outer skull, and scalp surfaces were segmented with the FSL toolbox using the individual T1 and T2 weighted MRI scans and converted into inner skull, outer skull, and scalp surfaces (Smith, 2002; Smith et al., 2004) that defined the boundaries for skin, skull, cerebrospinal fluid for each participant. The 3-D locations of the EEG electrodes at the surface of the head and the three major fiducials (nasion and left and right peri-auricular points) were digitized using a 3Space Fastrack 3-D digitizer (Polhemus, Colchester, VT) and co-registered with the anatomical scans. Visual areas were defined by fMRI retinotopic mapping (Tootell & Hadjikhani, 2001; Wade et al., 2002). hMT+ was identified using low contrast motion stimuli similar to those described by Huk & Heeger (2002). LOC was defined using a block-design fMRI localizer scan in which blocks of images depicting common objects alternated with blocks containing scrambled versions of the same objects. The stimuli were those used in a previous study (Kourtzi & Kanwisher, 2000). The area activated by these scans covers almost all regions (e.g., V4d, LOC, and LO+) that have previously been identified as lying within object-responsive LOC (Kourtzi & Kanwisher, 2000; Tootell & Hadjikhani, 2001).

To create a method that can be used widely across different EEG montages, we utilized the 10-05 system (Oostenveld & Praamstra, 2001) as a high resolution master montage with known fiducial locations. We aligned the previously measured and co-registered electrode locations to the 10-05 high resolution standard EEG system using an affine transformation based on 19 electrodes (Fp1, Fp2, Fz, F7, F3, C3, T7, P3, P7, Pz, O1, Oz, O2, P4, P8, T8, C4, F4, F8). We then combined the individually defined source space, surface boundaries and 3-D electrode locations with the MNE software package to estimate the electric field propagation with the standard Boundary Element Method (M. S. Hämäläinen & Sarvas, 1989). The resulting 50 forward models (one per individual) link the activity of the 20,484 cortical sources to the voltages at the surface of the scalp recorded by the standard 10-05 EEG system with an average reference.

To create EEG templates of functional brain areas, we pooled the forward model sources located within each of the 18 previously identified ROIs for each participant separately. We then projected the activity of each ROI to the scalp surface. The resulting scalp activity is different for each ROI and participant. However, because we created forward models with the same electrode layout for all participants, the scalp activity can be averaged across the 50 participants for each of the 18 ROIs. This averaged activity is what we term EEG templates. These templates can then be used to recover the sources of an observed EEG signal.

The advantage of using a 10-05 system with a high density of electrodes is that the templates can be fit to any EEG montage with the provided Matlab program (createCustomTemplates.m). We also provide EEG templates for EGI (Geodesic Sensor Net) montages which include electrodes located outside the 10-05 system. Most of the results are reported with a 128 electrodes EGI system but other montages are also compared in the results section.

### Recovering sources: calculating the inverse solution

Once EEG templates for each of the 18 ROIs are created, they can be used via linear regression to determine the brain sources responsible for the EEG activity recorded on the scalp (as long as the EEG montages and references match). It is well known that methods that fit more sources than sensors are “ill-posed” and require extra constraints and regularization (Baillet, Mosher, et al., 2001; Dale & Sereno, 1993; M. Hämäläinen et al., 1993; M. S. Hämäläinen & Ilmoniemi, 1994). Even for seemingly “well-posed” linear regression in our case (only 18 sources and 32-346 electrodes) solution regularization is crucial. This is because the forward model for EEG sources is typically not well-conditioned (because the eigenvalues of the forward model matrix rapidly diminish). The condition number (of the forward matrix) directly relates to the SNR that is required for perfect recovery of underlying sources. In our case, the condition number for the template forward models are on the order of 300. The SNR in typical ERP studies is lower than what would be required for this level of conditioning. Therefore, performing an un-regularized linear regression will result in excess noise being included in the solution (i.e., overfitting). In order to accurately fit the data, it is very important to appropriately regularize the solutions.

However, determining how much regularization to perform is a difficult open problem. Too much regularization results in underfitting the data and smoothing across real differences, while too little regularization results in overfitting and emphasizing noise which enhances spurious patterns. Choosing the amount of regularization can be done subjectively by the experimenter or by using an algorithm. The present paper uses two of the popular algorithms for choosing regularization amounts: the Generalized Cross Validation (GCV) and the L-curve method (Hansen, 1992; Hansen & O’Leary, 1993). We used the Regularization Tools Toolbox (Hansen, 1994) and made modifications to extend the L-curve and GCV functions to be applicable to data with multiple samples (e.g. over time for an ERP). Both the GCV and L-curve methods work by generating proxy estimations of over/under-fitting and optimizing this tradeoff. GCV methods have been very useful in the past for regularizing EEG source localization solutions in individual participants (Cottereau et al., 2015). However, we found that with the template method applied to average data, the GCV can fail to provide an appropriate regularization value. This is due to the GCV’s assumption that residual error for left out data will be independent (uncorrelated) across electrodes. Indeed, the residual error across electrodes can be highly correlated with the template method, biasing the GCV estimate of fitting. In these circumstances, the L-curve method is better suited than a GCV regularization since it is more robust to the presence of correlated errors (Hansen & O’Leary, 1993).

Because both the L-curve and GCV methods rely on strong assumptions to provide approximate regularization parameters they are not guaranteed to provide an optimal value. Optimal regularization is still a fundamental and challenging problem. Active research is ongoing for developing methods that would provide a better regularization estimation and future results may improve on this algorithm. When simulating data we have access to the true sources of the signal, it is thus possible to choose a regularization parameter that best optimizes the error (based on computing the Mean Square Error) instead of relying on the L-curve or GCV regularization. In order to evaluate the best-case scenario that may be achievable in the future with improved regularization methods, we also performed source localization using this best possible regularization parameter that we call the *template-optimal* method.

### Comparison with other source localization methods

Traditional source localization methods cannot be directly compared to the proposed method. Because our goal is to recover the signal from functional sources, we consider only 18 candidate sources in our procedure whereas traditional methods recover anatomical sources among candidates distributed throughout the whole brain. However, it is important to validate our approach by comparing it to other source localization methods that are based on each individual head model. We therefore compared our method to those that have been previously used for fMRI ROI-informed source localization (Cottereau et al., 2015). We localized sources from each individual separately and pooled the identified brain sources within each ROI (sources outside the ROIs are ignored). The recovered source activity within each ROI is then averaged across participants and can be compared with the results of the template method (Fig.1). While this *individual* method considers any point of the cortex as a potential source, we also performed source localization considering only the brain points located within the ROIs. This *individual-subset* method is more focused and has assumptions comparable to the template method. In addition, for simulated data, we also compared the template method with an *individual-oracle* method. Because we have perfect knowledge of the true sources of the signal, it is possible to use the same data for simulating and retrieving the sources. In our case, we use the same 18 ROIs of each individual participants to simulate and retrieve the data. This *individual-oracle* method gives the upper bound of source recoverability.

**Figure 1.**
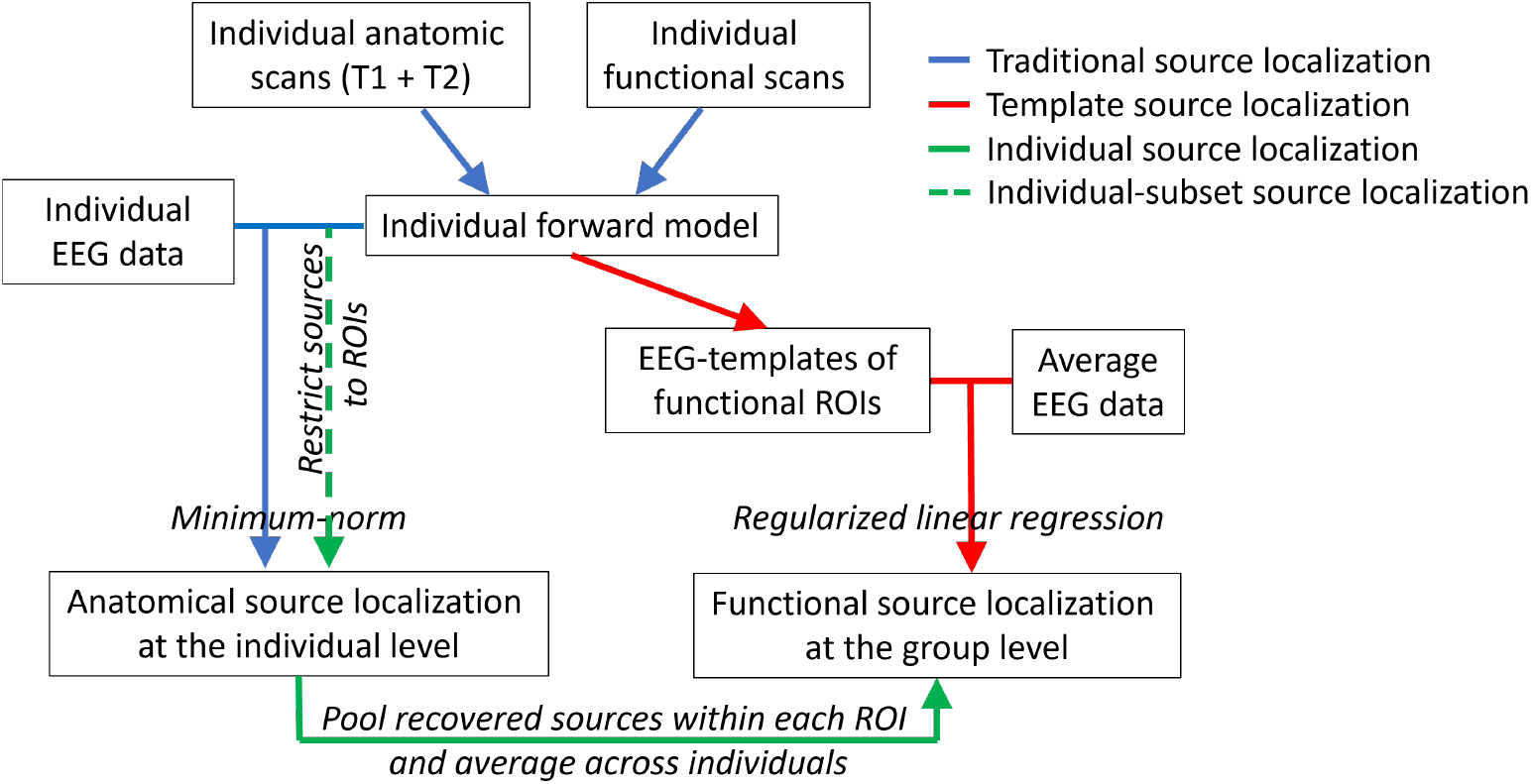
Analysis pipeline for different source localization methods. The template method uses EEG templates (representing typical average scalp activity for a set of ROIs) to retrieve functional EEG sources at the group level. Traditional source localization methods use individual scans to retrieve anatomical EEG sources at the individual level. To retrieve functional sources at the group level, we pooled and averaged the sources; a method that we call “individual source localization” or “individual-subset source localization” when we restricted the potential sources to be only in the ROIs (see text for more details).

In the case of all these source localization methods, either of the L-curve or GCV method could be used for determining the regularization parameter. We tried both and found that source localization results were generally better when using the GCV at lower SNR. We thus report the results of the individual-based methods using GCV regularization (results using the L-curve can be found in Supplementary Material).

### Data simulations

To test the template method, we simulated scalp EEG data with known brain sources and attempted to retrieve them while varying different parameters such as the amount of noise in the data and the number of participants. The first set of simulations consisted of testing the amount of similarity between templates or, in other words, assessing the crosstalk (leakage) between ROIs. This is important for determining the type of errors the template method commits (i.e., which ROIs are confused with one another) and whether these errors reflect realistic errors (ROIs that are anatomically close are more confusable than anatomically far ROIs). For this, we simulated the activity of one ROI using the forward models of a set of 50 participants. This resulted in different EEG scalp signals for each participant to which Gaussian white noise was added to all electrodes to obtain a signal-to-noise ratio (SNR) of 200. The SNR was defined as the ratio between the variance of the signal and the variance of the noise across time. The simulated signal was then recovered using the template method and other source localization procedures described in the previous section.

In another set of simulations, we tested source localization performance when a pair of bilateral brain sources (two sources in each hemisphere) were active using a more realistic ERP-like signal of various amplitudes. The pair of sources was chosen as being either easily distinguishable, V1 and hMT+, or difficult to separate, V2v and V4 (there is a considerable amount of crosstalk between V2v, V4 and V3v, which separates V2v and V4). For each simulation, a source signal was created over time with a baseline activity from −45 to 0 ms, a bilateral V1 (or V2v) response from 0 to 45 ms, a bilateral hMT+ (or V4) response from 46 to 90 ms and a combined V1 and hMT+ (or V2v and V4) response from 91 to 135ms (Fig. S1). The source signal was created with a random amplitude between 1 and 10 and had an ERP-like shape that was common across all participants but different for each simulation. From these brain sources, we simulated a scalp response using the forward model of the participants that were included in the simulation. The participants (N=2, 8, 20 or 50) were randomly chosen, with replacement, for each simulation. Gaussian noise was added to all electrodes of the EEG signal corresponding to SNR levels of 0.1, 1, 10, 200, 10 000. These SNR levels extend beyond the SNR level observed in recorded data (an SNR of 0.1 corresponds to an activity with 10 times more noise than signal, while an SNR of 10 000 corresponds to 10 000 times more signal than noise). Given that the forward models and noise are not common across individuals, the EEG scalp data will be different across individuals. Note that adding noise to the electrodes is the same as pre-whitening the signal. For each parameter combination (pair of sources tested, number of participants, SNR level), we ran 30 simulations.

To test different EEG montages (with 32, 64, 128 and 256 electrode EGI systems), we used similar ERP-like simulations but instead of using V1-hMT+ or V2V-V4, the two active pairs of bilateral areas were randomly chosen. We also tested the effect of the number of simultaneously active ROIs on source localization performance for a 128 electrodes EGI montage by simulating the activity of each bilateral ROI as active (1) or non-active (0) with up to 9 bilateral ROIs active simultaneously (i.e. 18 visual ROIs).

### Application to real data

The difficulty in testing source location methods when using real data is that we do not know the true sources, so it is not possible to establish whether a method is better or worse than another. Here we applied our template method to a dataset that had been used to test functional source localization methods (Lim et al., 2017). The dataset consists of EEG recordings of 9 participants who viewed dynamic Random-Dot-Kinematograms (RDK). The RDK alternated every 500 ms (at 1 Hz) between incoherent and coherent rotary motion (alternating between clockwise and counterclockwise direction to reduce the effects of motion adaptation). EEG responses at 1 Hz (and its harmonics at 2Hz, 3Hz, etc.) reflect changes in global motion and can be interpreted as arising from areas that can discriminate between coherent and incoherent motion (Norcia et al., 2015). Given previous fMRI and MEG results contrasting these two types of motion, we expect sources to be present in V3A and hMT+ (Aspell et al., 2005; Costagli et al., 2014; Händel et al., 2007; Helfrich et al., 2013; Lim et al., 2017; Rees et al., 2000; Rina et al., 2022).

The EEG data was collected using a 128 electrodes EGI system. The data was referenced to the average signal and matched with the reference of the templates. The average reference was chosen because it avoids having the reference channel exert extra influence on solutions (Hu et al., 2018; Yao et al., 2019). This is important for both the presently proposed template method and other localization methods.

Source localization was conducted on the first five harmonics of the signal (1, 2, 3, 4 and 5Hz) using the template, individual and individual-subset methods. Because the same dataset was previously analyzed using a group-lasso procedure (Lim et al., 2017), we also reproduced that analysis. The source localization was repeated 500 times with a different sample of participants. Using this bootstrap distribution, we computed a 95% confidence interval and tested whether a given ROI was active at any time point (comparison with null activity using alpha=0.05). In more traditional paradigms comparing two conditions with ERPs, we advise users to perform permutation tests between the two conditions to test for significant differences. For plotting purposes, the retrieved activity was normalized across sources by the maximum retrieved activity across time and ROIs.

### Evaluating source localization results for simulated data

Source localization methods can be evaluated in a wide variety of ways. In this study we utilized three metrics that emphasize different aspects of source localization error. Each of these metrics is calculated at each time point on the recovered signal then averaged across time.

<u>Area Under the ROC Curve (AUC):</u> quantifies how well the source estimation procedure discriminates between active and non-active sources. The ROC curve compares the sensitivity (true positivity rate) and the specificity (true negativity rate) at different activity thresholds (criterion in signal detection terms). AUC varies from 0.5 (random classification between active and non-active source) to 1 (100% correct classification for all threshold levels, no false positives or false negatives).

<u>Relative energy:</u> specifies the amount of energy recovered in the active sources by quantifying the amount of leakage of energy outside the correct solution. It is the ratio between the normalized estimated activity contained in the true active sources and the normalized estimated activity in all possible sources. A perfect estimation with no leakage results in a value of 1.

<u>Normalized Mean Squared Error (MSE):</u> measures how close the recovered amplitude is to the simulated source amplitude. It is the average of the square of the difference between actual and estimated EEG activity normalized across sources at each time point. MSE is always positive and decreases as the error approaches 0. It reflects the fit to the ground-truth signal.

## Results

### EEG templates (distribution of scalp activity) for the 18 ROIs

Our template-based source localization approach relies on using EEG topographies that represent the activity of intracranial brain sources. We thus modelled the EEG activity of the 18 ROIs that we considered (see Methods) for 50 participants. The resulting individual topographies for a given ROI show some similarities but also clear differences, illustrating cross-participants variability (some examples can be found in Fig.S2). This variability primarily reflects anatomical and functional brain differences between participants. A minimal amount could also be explained by variations in the processing steps for creating the forward models (for defining the source space, surfaces boundaries or for ROI localization). Such variations are expected and part of any analysis so they are a good representation of the type of data that would be recorded by researchers.

The average scalp activity for each ROI is illustrated in Fig.2 for a 128-channels EGI system. The topographies of the EEG templates for a 10-05 system with 346 electrodes (Oostenveld & Praamstra, 2001) and common montages, including the ones tested in this study with 32, 64 and 256 electrodes can be found in Fig.S3. Apart from showing the EEG templates that we use for the source localization in our simulations; these figures are important as they depict the EEG topography to expect when specific brain areas are active. It is worth noticing that for example, when V1 in the left hemisphere is active (which would be expected for a stimulus presented in the right visual field), it results in a maximum scalp response in the opposite hemisphere (posterior-right of the head). This paradoxical lateralization of the EEG response (ipsilateral to the visual hemifield that is stimulated) has been documented almost 50 years ago (Barrett et al., 1976). The V1 template demonstrates that this paradoxical activation is observed when averaging over a large sample of participants and is the direct consequence of brain anatomy.

**Figure 2.**
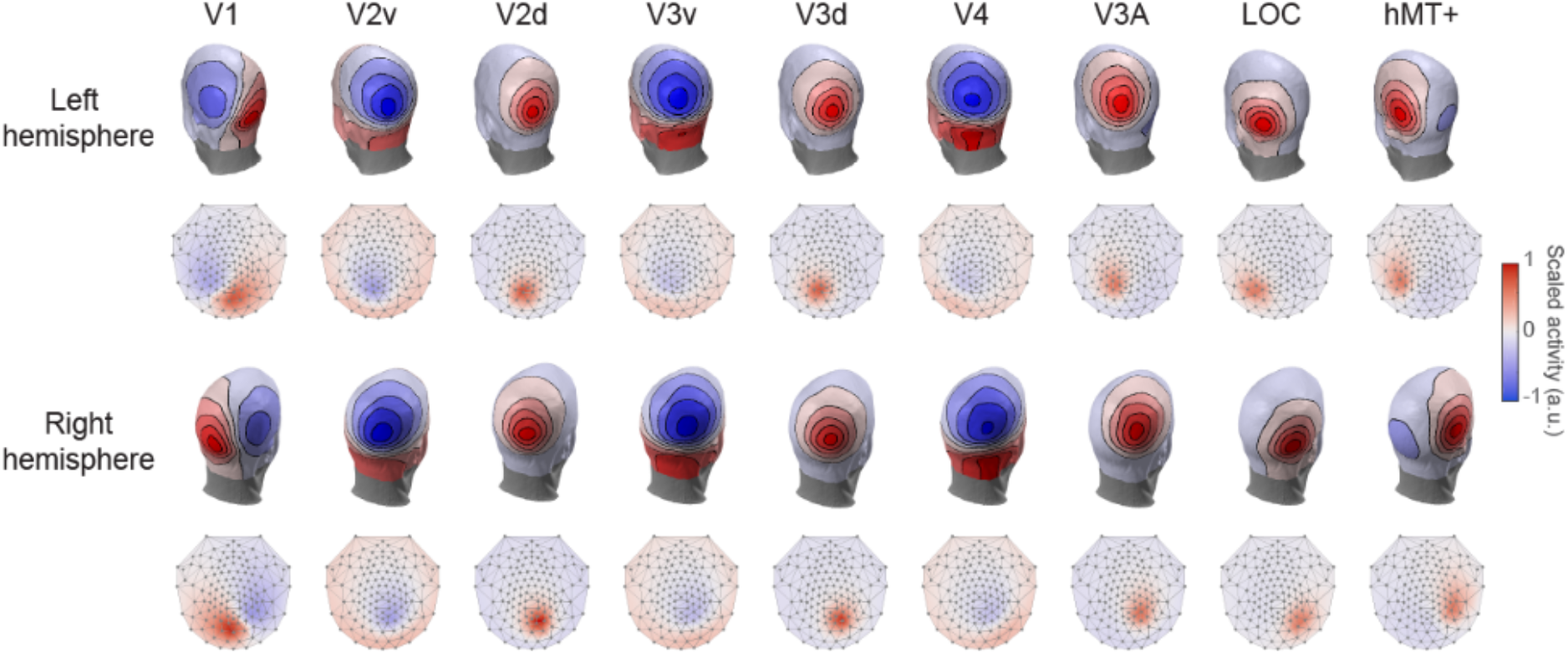
EEG templates using a 128 electrodes EGI montage represented on a 3-D and 2-D head layout. The sources were first defined for each individual participants using anatomical and functional scans. Using individual forward model, the activity of each source was then projected onto the surface of the scalp and averaged across participants. The intensity of the color indicates the amplitude of positive (red) and negative (blue) activity.

The activity in ROIs such as V2D, V3D, V3A, LOC and hMT+ result in a more focal positive signal on the scalp. Note that the scalp activity resulting from hMT+ is more anterior than what might be expected. Indeed, the activity from V3A could easily be mistaken as that arising from hMT+. This observation highlights the utility of such templates: in addition to being instructive, having a good idea of the scalp activity that results from different brain areas can help interpreting EEG results. Moreover, because different electrodes represent the activity of certain ROIs, such a figure can be used as an a priori method to decide which electrodes to pool or keep separated when analyzing EEG data. Additionally, these can provide an alternative to fiducials to compare responses across different recording montages.

The resulting scalp activity of some ROIs are similar with each other (e.g., V2D and V3D), while the activity of other ROIs (e.g., hMT+) show distinct topographies (Fig.2). High similarity between EEG templates for different ROIs can increase the confusion in retrieving the true source of a signal. We estimated this confusion by computing the amount of crosstalk between ROIs, which represents the amount of activity arising from an active ROI that is attributed (leaks) to other non-active ROIs. Crosstalk is higher within than between hemispheres, within early ventral areas and within early dorsal areas (Fig.3; results for other source localization methods can be found in Fig.S4). This reflects the anatomical and functional similarities between ROIs, as one would expect from any realistic individual-based source localization.

**Figure 3.**
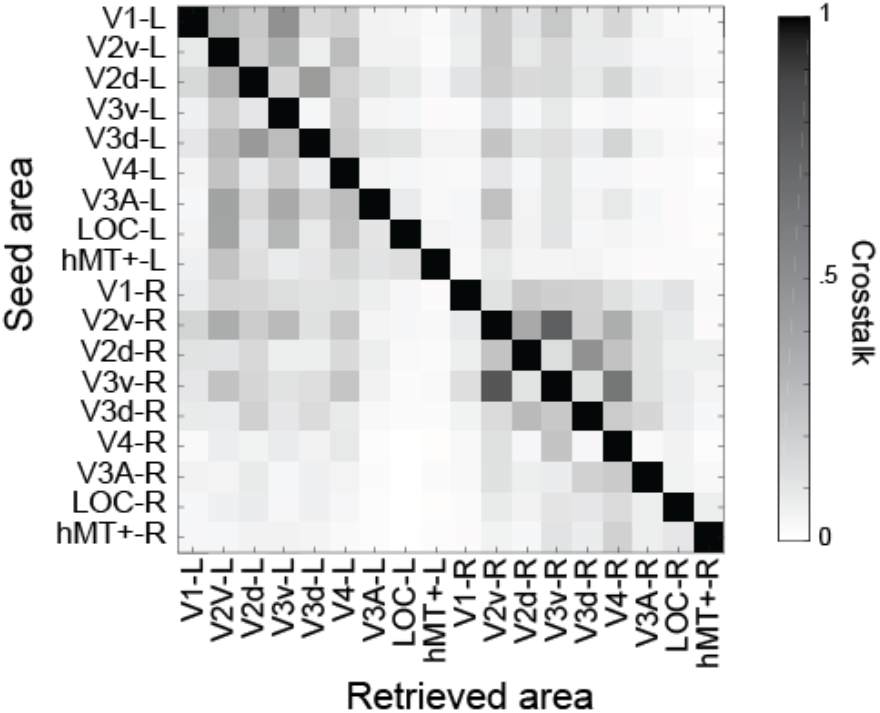
Crosstalk (leakage) between ROIs using the template method. The amount of crosstalk (normalized for each ROI; per row) was calculated for an EEG signal simulated using a 128 EGI montage with an SNR of 200 and averaged across 50 individuals. In an ideal although implausible system, only the diagonal will be dark with the rest being white (zero crosstalk).

The EEG templates were created from the modelled scalp activity of 50 participants that were then averaged. Although one can always add more participants to create these templates, it is interesting to consider that the variability in these templates is inversely proportional to the number of participants (Fig.S5). Such exponential decay slope (1/N for variance and 1/sqrt(N) for standard deviation; where N represents the number of participants) is expected when averaging random samples. For each additional sample, there is diminishing returns such that for reducing variance in half, the number of participants needs to quadruple. Thus, it would require a large number of participants to make a marginal improvement in the population sampling error present in the current study. Therefore, these EEG templates can be confidently considered a good representation of each ROI activity.

### Simulation results

#### Comparison between average and individual-based source localization

We simulated EEG activity for a 128 electrodes EGI system from a signal originating in V1 and hMT+, or in V2v and V4 (see Methods). The simulated EEG was created with different SNR which corresponds to different level of noise in the signal but can also be taken as a proxy for the number of trials averaged within one experimental condition (i.e., as the number of trials increases, SNR also increases). While the extreme SNR values of 0.1 and 10000 are unrealistic and reflect lower/upper bounds, an SNR between 10 and 200 reflect typical experimental conditions where ERP results are averaged across hundreds of trials (for ERP examples with different levels of SNR, see Fig.S6).

Using the source localization procedures detailed in the method section and summarized in Fig.1, we recovered, time point by time point, the activity of the brain areas generating the simulated EEG response (Fig.S1). The source localization performance was assessed using three metrics (see Methods) for varying numbers of participants and levels of SNR. The template method clearly does very well, with performance close to the other procedures that use individual forward models (Fig.4A). As expected, performance improves with higher SNR and more participants. Performance is relatively poor when the data of only 2 participants is simulated but the template method does equally well for recovering activity in V1 and hMT+ with 8, 20 and 50 participants. The metrics improve with higher SNR until they reach an asymptote. Source localization results for the template method are slightly lower than the individual method, while the individual-subset method shows better results. Indeed, the metrics for the individual-subset method are almost as high as the ones for the individual-oracle method, demonstrating that V1 and hMT+ are easy to separate at an individual level, especially with high SNR.

**Figure 4.**
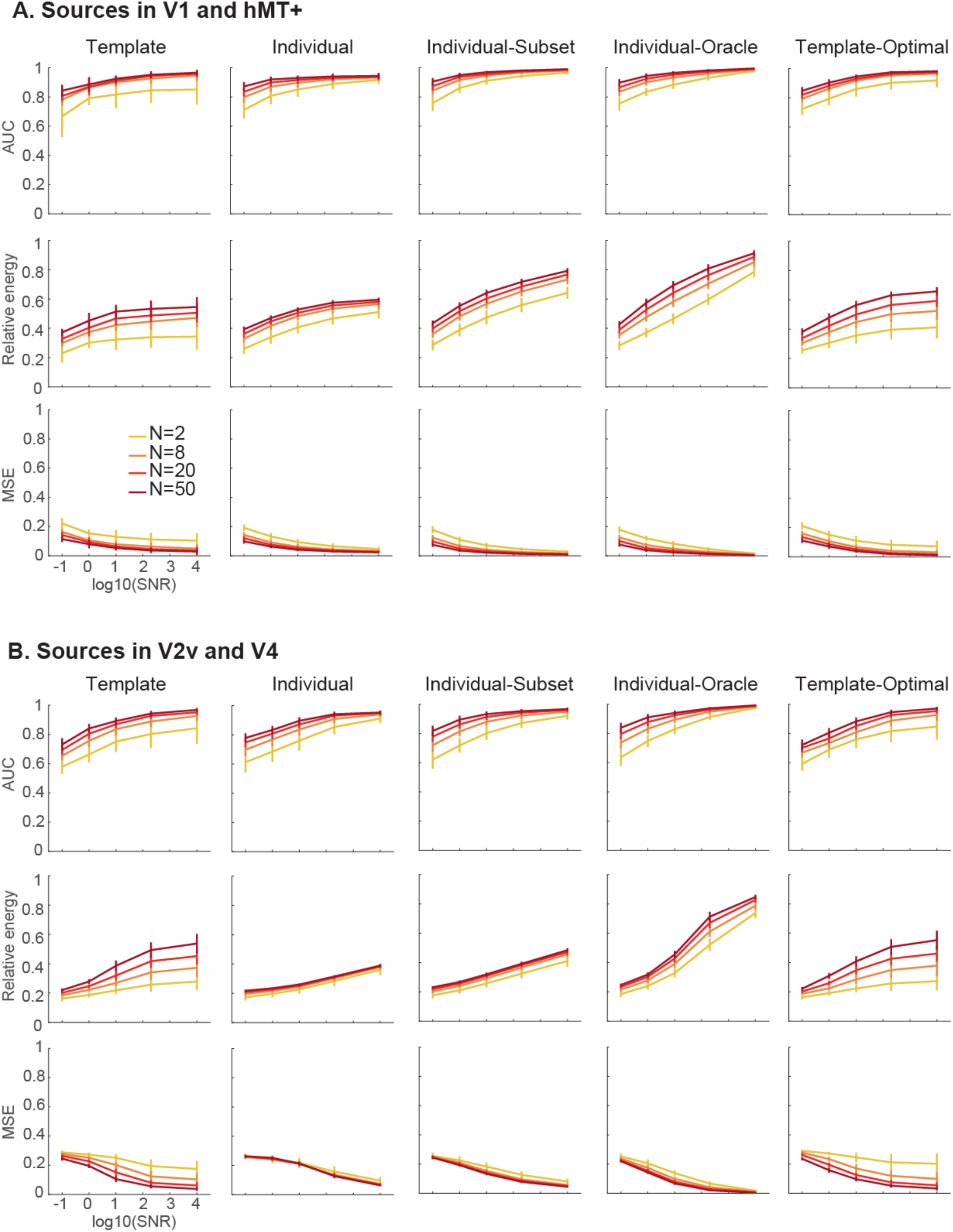
Source localization performance for a signal simulated with 2, 8, 20 or 50 participants, a 128 electrodes EGI montage and different SNR levels. The Area Under the Curve (AUC) quantifies whether the true sources have been recovered; the Relative Energy measure quantifies the amount of leakage; the normalized Mean Squared Error (MSE) quantifies the difference between the true and the recovered signal. Sources are simulated bilaterally in V1 and hMT+ (A) or in V2v and V4 (B). Error bars represent standard deviation.

On the other hand, and as expected, source localization results for all methods are lower when attempting to localizing activity generated in V2v and V4 (Fig.4B). These two ROIs and V3v are known to exhibit substantial crosstalk. This can be seen in Fig.3 and by their template resemblance in Fig.2. Hence, this situation is challenging for source localization. Nevertheless, when V2v and V4 are active, the template method does well, if not better, than other methods, particularly when a large number of participants are simulated (although, as expected, still not as well as the individual-oracle method). This directly illustrates the advantage of averaging across participants.

When a pair of sources are easy to separate, averaging the EEG across participants does not improve source localization much. However, for sources that are close to each other, averaging the EEG scalp activity leads to a better characterization of the signal common to all participants, making it easier to separate them from each other and from other potential sources. Thus, despite not being tailored to each participant, the template source localization method that we have developed recovers active functional ROIs equally well in both easy and difficult situations.

Compared to the template-optimal method (which uses the best possible regularization parameter, see Methods), the template method is slightly less accurate when simulating activity in V1-hMT+ but not very different when simulating activity in V2v-V4. This demonstrates that the regularization parameter that we use (based on L-curve) is appropriate in various circumstances.

#### Comparison between EEG montages

Previous studies have shown that low spatial sampling can lead to incorrect source localization and that higher spatial sampling increases source localization precision (Brodbeck et al., 2011; Srinivasan et al., 1998; Staljanssens et al., 2017; G. Wang et al., 2011). Similarly, the success of our new source localization method might also depend on how precisely the EEG templates are defined: with different number of electrodes, the topographies of the templates and of the EEG dataset will be more or less precise. We thus tested our template method using montages that included 32, 64, 128 or 256 scalp electrodes (note that the system itself, EGI, Biosemi or others, would not affect the results of our method; the EEG templates can be used with any EEG system as long as the templates and the data on which source localization is applied have matching montages). Localization performance improves with more electrodes but is still reasonably good for fewer electrodes (Fig.5A). This pattern is also observed with methods using individual forward models (Fig.S7). This demonstrates the validity of the template method for a large range of EEG montages.

**Figure 5.**
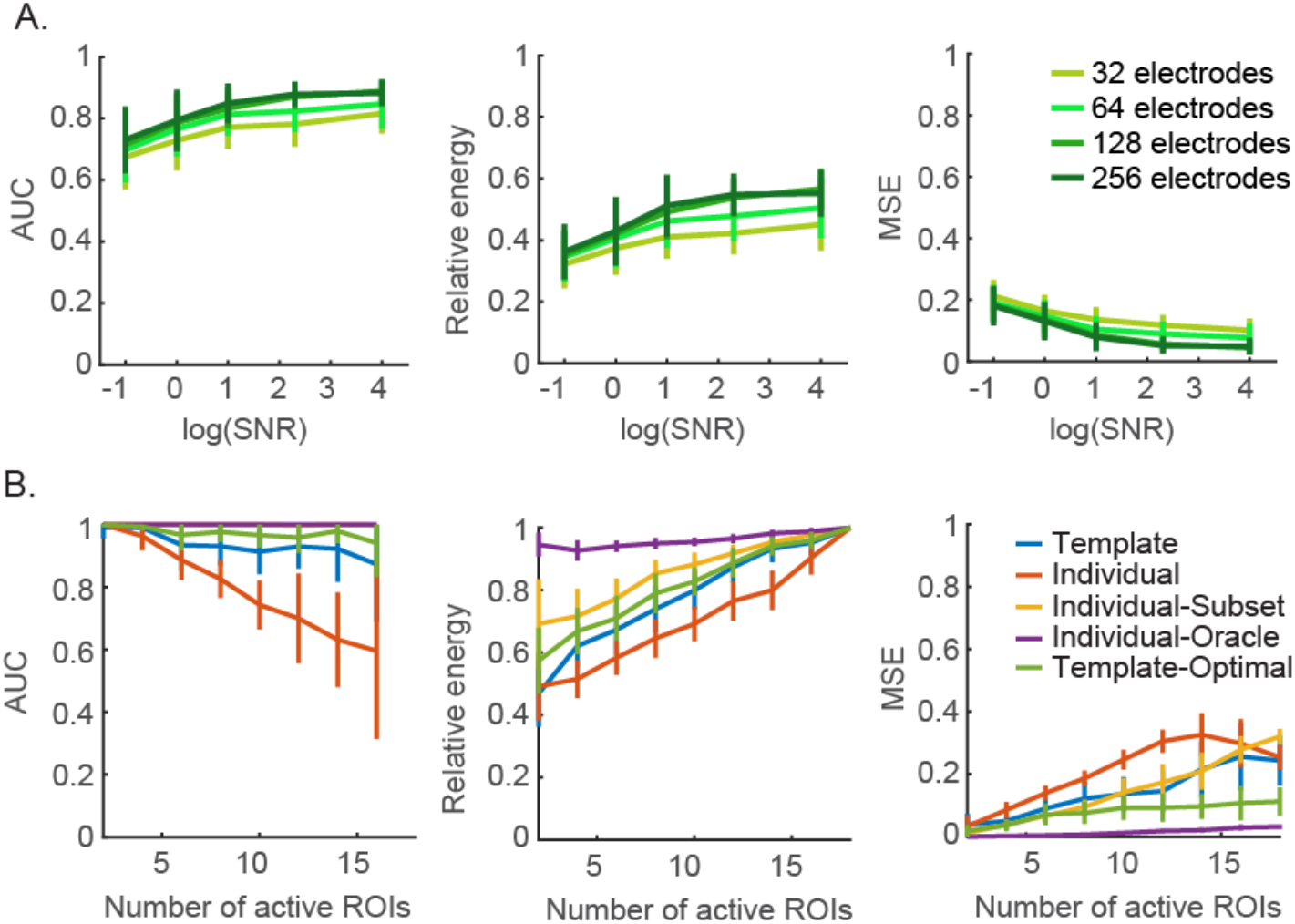
A. Performance of the template-based source localization method for EGI montages with 32, 64, 128 and 256 electrodes. Brain sources are recovered from the simulation of two randomly chosen bilateral ROIs. B. Source localization performance using different methods for retrieving 2 to 18 simultaneously active sources. Sources are simulated bilaterally for a 128 electrodes EGI system.

The pattern seen in these simulations demonstrates that data quality matters and can be somewhat traded-off with fewer electrodes. At the lower end of SNR, larger improvement in source localization performance can be attained from improving SNR rather than from adding more electrodes. However, at the upper end, increasing SNR provides diminishing returns and the enhancement in source localization accuracy provided by more electrodes is more important.

#### Number of active ROIs

In addition to simulating the activity of a pair of ROIs, we also tested the reliability of the template method for an increasing number of ROIs active simultaneously (Fig.5B). The AUC for the template method decreases with an increasing number of ROIs but stays reasonably high (around 0.9). Not surprisingly given how relative energy is calculated (ratio between the energy in active ROIs and the energy in non-active ROIs), it increases with increasing number of simultaneously active ROIs. This is also the case for the MSE. The interesting comparison here is across the different source localization methods. The individual method does poorly when multiple sources are simultaneously active: the classification between active and non-active sources (AUC) is below 0.8 and almost at random (0.5) with 16 sources active simultaneously. The template method’s AUC and relative energy are higher than the individual method but lower than the individual-subset method. MSE is similar for both the template and the individual-subset method, although with a better regularization term, the MSE for the template-optimal method stays lower. Thus, the accuracy (AUC) of source localization using the template method decreases a bit with increasing number of simultaneous active sources but its performance is still high. The effect of the number of active sources on source localization results is rarely tested but we show here that our method performs well regardless of how many sources are active simultaneously.

### Real-data results

We tested the template method on a real EEG dataset collected while participants viewed RDK alternating every 500 ms between coherent and incoherent motion. Because we do not have access to the true active sources, we cannot compute goodness of fit measures such as the AUC, relative energy and MSE. We can only compare the results of the template method with other individual-based methods and with previous studies using a similar paradigm. We also compared the results with the ones from Lim et al. (2017) who analyzed the same dataset with the group-lasso method. With that method, the source localization is performed using each individual’s forward model but the information from all participants is used to ensure that the selected sources are in agreement across all participants.

Source localization results are very different between the source localization methods (Fig.6). While most ROIs are active when using the individual method, very few are when using the individual-subset method. Although the individual-subset method had better results in the simulations, here the retrieved sources are inconsistent between the left and right hemispheres. On the contrary, when using the template or the group-lasso method for the source localization, a couple of ROIs, V3A and hMT+, are clearly active throughout the presentation of the stimuli and the waveforms are very similar. V3A and hMT+ are the two areas that are commonly found to be sensitive to motion coherence (Aspell et al., 2005; Braddick et al., 2001; Händel et al., 2007; Helfrich et al., 2013; Rees et al., 2000; Rina et al., 2022) so these results demonstrate the validity of our method when used on real data. A similar waveform is also observed in V4 which is consistent with an fMRI study that showed that V4 responds to transient increments and decrements in the level of motion coherence (Costagli et al., 2014).

**Figure 6.**
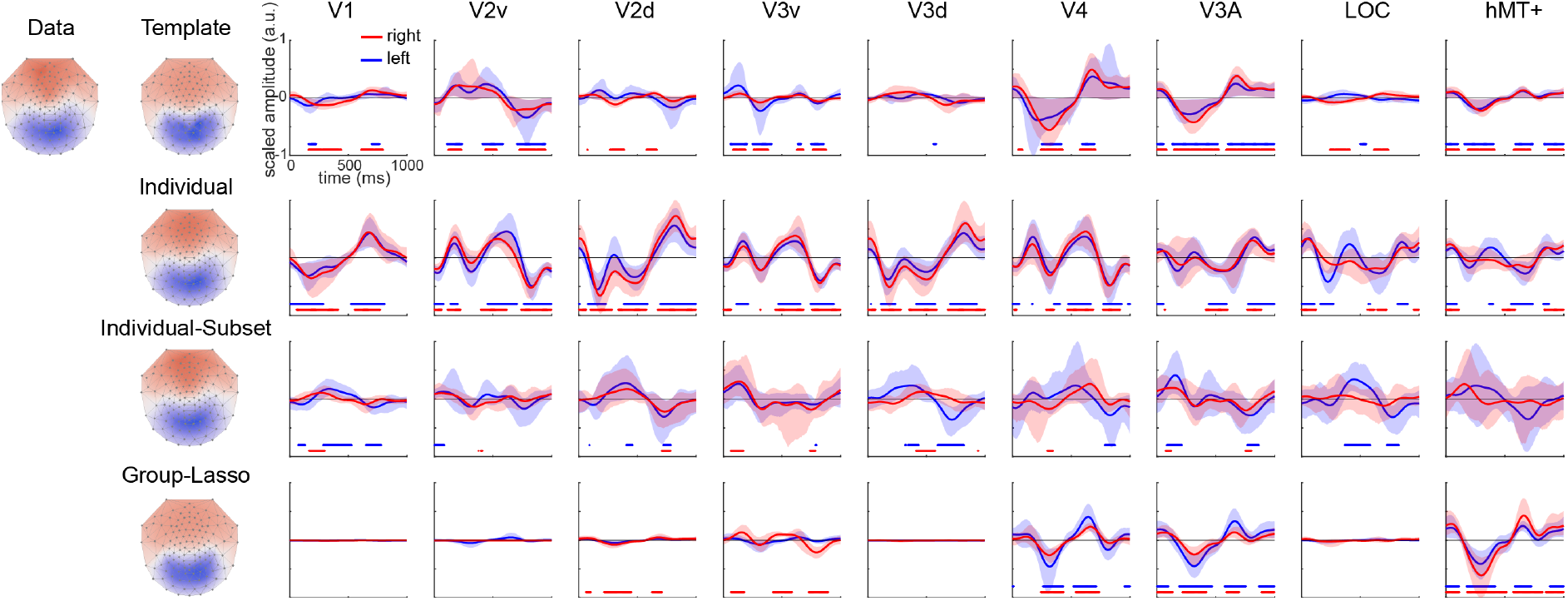
Source localization on a real EEG dataset collected while 9 participants viewed coherent and incoherent motion alternating every 500 ms. Observed and reconstructed topographies at 250 ms are shown (left) along with the time course of retrieved sources in different ROIs using various source localization methods (right). The average ROI activity computed over 500 bootstraps is represented in blue for the left hemisphere and in red for the right hemisphere with the shaded area representing 95% confidence interval. The stimulus was viewed centrally, and we would thus expect the brain activity to be symmetrical between the left and right hemispheres. Active ROIs (bootstrapped distribution different from 0 at a threshold of p<0.05) are indicated by blue and red dots (for left and right hemispheres respectively) at the bottom of each plot.

We also observe some activity in early ventral areas V2 and V3 when using the template source localization method. Interestingly the most active area among those, V2v, shows a waveform that is inverted compared to V3A and hMT+. This indicates that the recovered activity is not due to crosstalk or leakage from V3A and hMT+. Previous studies have also reported activation of areas located on the posterior ventral surface of the occipital lobe (Braddick et al., 2001; Händel et al., 2007; Rees et al., 2000; Rina et al., 2022). Interestingly, some of them (Braddick et al., 2001; Rina et al., 2022) found that responses in early visual areas (V1/V2) decreased as coherent motion strength increased, which is the opposite of what is observed for areas hMT+ and V3A. The results of the template method showing inverse activity in early ventral occipital areas compared to hMT+ and V3A is thus consistent with these reports.

It is interesting to note that, as is usually found in EEG studies, the data shows negative activity in posterior electrodes together with a positive activity in frontal electrodes (here illustrated at 250 ms in Fig.6). Despite no specific ROI template showing such pattern, all source localization methods retrieve and account for this topography based on only sources in visual areas. This illustrates how the combination of multiple active sources can create unexpected EEG topographies. That is, activation in frontal electrodes need not depict activity in the frontal cortex but can arise through combination of activities in occipital areas.

Analysis on this real-world dataset shows that the template method clearly surpasses the methods that use individual head models that do not use or pool information from the group. Further, the source localization results are almost uninterpretable using the individual and individual-subset method. On the other hand, results from the template method echo previous ones from fMRI and MEG experiments. The superiority of the template method may be explained by differences in SNR. When using individual head models, source localization is applied to each participant’s data. This data has a lower SNR than the data averaged across participants. The template method on the other hand is based squarely on using averages to improve source localization.

## Discussion

We present a new EEG source localization method that is simple, efficient, inexpensive, and rapid. This method relies on functionally defined EEG templates that we created by modelling the average EEG scalp response for each of a set of 18 functional ROIs. With these templates, we can then estimate the contribution of each ROI to the recorded EEG signal using a regularized linear regression. We tested this template method in an extensive set of simulations and the results show that it is as efficient as other methods that use individual forward models. With just 8 participants (and a realistic SNR level of around 200), the sources at the origin of the EEG signal are correctly identified. We also show that the source localization accuracy of the template method is reasonably high even with montages of a small number of electrodes and with a large number of simultaneously active ROIs. Performance of the template method is particularly impressive when tested on a real dataset. It retrieves brain sources in a limited set of ROIs and these ROIs correspond to the ones identified in previous reports (Aspell et al., 2005; Braddick et al., 2001; Costagli et al., 2014; Händel et al., 2007; Helfrich et al., 2013; Lim et al., 2017; Rees et al., 2000; Rina et al., 2022).

Contrary to other source localization approaches, the template method uses the averaged EEG across participants to retrieve the brain sources. Thanks to this, the SNR of the data is high and results in accurate localization of the sources. In fact, applying source localization on an individual basis, as done with traditional source localization analyses, amplifies the noise in the data which can lead to retrieving erroneous sources. Source localization methods using individual head models suffer from the difficulty of needing a principled way to average across participants since estimated anatomical sources do not match across participants. Here, we use what is common across participants to improve source localization at the group level and reduce cross-participant variability originating from functional and anatomical brain differences. This also means that our approach is best suited for group-level analysis.

In all source localization methods, the amount of retrieved activity is represented in arbitrary units and comparing this amount across brain sources is always difficult. Indeed, the amount of scalp activity of an ROI depends on the size of the ROI. For example, because V1 is bigger than other ROIs, it will have a stronger scalp response, which is what we observe in the EEG templates (Fig.2). To take into account the size of the ROIs, the scalp activity can be normalized such that the total power across electrodes for each ROI is equal to one. However, in that case, the activity in smaller ROIs will be overestimated and that in larger ROIs will be underestimated. We have chosen not to normalize the EEG templates as we think that they are a better representation of the brain activity. However, it is important to keep in mind that comparing the amount of activity between ROIs is difficult. It is more appropriate to determine whether an ROI is active or not (just as the simulation results also show, AUC is high but the relative energy performance is lower). Similarly, when comparing two conditions, it would be more advisable to focus on differences in activity within the same ROI across two or more conditions than across ROIs.

Beyond the source localization aspect, the scalp representation of the activity of different brain sources, the templates, are useful for understanding EEG topographies in general. It can be fairly difficult to interpret scalp topographies. This is because the forward solution for how a specific source in the brain appears on the scalp is nonlinear and the brain surface is convoluted and complicated. A good demonstration of this is the paradoxical lateralization of EEG responses to visual stimulation where the maximal responses can appear on the “wrong” side of the head compared to the known source of the signal (Barrett 1976). This happens due to the specific shape of the cortex causing the sources to point in unexpected (but measurable) directions. This is also the underlying reason for the failure of the cruciform “orientation flip” model to isolate V1 components (Ales et al., 2010; Ales, Yates, et al., 2013). Similarly, we observe a positive frontal scalp response in the real data even though no individual template shows such activity. Instead, this topography is the result of the simultaneous activation of multiple posterior brain sources. By providing a set of templates for how functionally defined brain regions appear on the scalp, we can help develop a qualitative understanding of scalp distributions that can aid researchers in understanding and interpreting their data.

### Advantages

The proposed method has several advantages. The main one is that since it is based on EEG templates, it does not rely on any individually defined MRI or fMRI data. This implies that the method is substantially cheaper and also quicker since no additional scanning time or extensive processing of MRI or fMRI data is required. The second main advantage of the template method is that the results are interpreted in terms of functional ROIs, not as anatomical locations in the cortex. This is important since most studies, aside from certain clinical ones, use source localization methods to identify the functional brain areas contributing to the signal and are not inherently interested in specific anatomical sources. In addition, because the same ROIs are often defined in other neuroimaging studies, the results from the template method can be compared to MEG, fMRI or single-unit studies.

Another practical advantage of this method is that it is easy to use. It relies on a small set of functions (programmed in Matlab but easily transferable to other programming languages) that can be downloaded from https://github.com/aleslab/eegSourceTemplateMatching. We provide the EEG templates for EGI (Geodesic Sensor Net) and for a 10-05 system (346 electrodes) which can be subsampled from to match a different montage and/or electrode reference using a custom-built function that automates it (createCustomTemplates). Code that interfaces with EEGLAB (Delorme & Makeig, 2004) and FieldTrip (Oostenveld et al., 2011) is available, making this method accessible to a wide userbase.

A final advantage of the template method is that the solution to the linear regression (the contribution of each ROI) does not depend on the reference of the montage. Topographies and ERPs vary depending on the specific reference used for EEG analysis (Joyce & Rossion, 2005; Luck & Kappenman, 2012). With our method, regardless of the reference (as long as it is the same between the EEG templates and the collected data), the solutions to the regression will be the same. This thus increases the potential for comparisons across studies. However, it is important to note that in general the referencing scheme influences the accuracy of source recovery and an average reference scheme should be used for best results (Yao et al., 2019).

### Limitations & future directions

The main limitation of the method that we present here comes from its strength. Because it relies on a set of scalp activity templates, it considers only sources from a restricted number of ROIs for which a template is provided. Here, we created 18 EEG templates which are all in visual areas and thus expect no activity in other parts of the brain. For visual experiments, most of the brain response comes from the visual cortex so we can assume that brain sources are within the visual ROIs. Other brain areas might be involved in later stages of processes (i.e., for decision making) but the main interest of vision research studies is to reveal the neural mechanisms involved in visual perception, that is, within the 18 visual ROIs. Thus, even with a limited number of EEG templates, the template method can be used in a large range of visual experiments.

Importantly, following the same method as described here, new templates for other ROIs can be created. Although the current set covers a large part of the visual system, areas such as V6 (Cardin et al., 2012), KO (Tyler et al., 2006) or category selective brain regions responding preferentially to faces (FFA, Kanwisher et al., 1997), places (PPA, Aguirre et al., 1998; Epstein & Kanwisher, 1998), bodies (Downing et al., 2001; Peelen & Downing, 2005) and words (Cohen et al., 2000) could be localized with fMRI scans and their activity simulated at the surface of the scalp. Similarly, non-visual brain areas defined anatomically or functionally (for example, areas in the auditory cortex; Barton et al., 2012) could be added. This will extend the reach of the template source localization method that we developed to other research fields. In the same vein, one can create MEG templates to apply on MEG data to further ease the comparison between neuroimaging studies.

A second limitation of the template method is that it assumes that the average of multiple participants is a good representation of the group. This applies to the normal population, but this assumption might not be valid in clinical settings. For example, it might not be accurate to assume that brain processes are the same in stroke or epileptic patients. However, future research could use these templates to interpret differences in EEG topographies between individuals or between an individual and the templates. Similarly, the EEG templates might not be representative of the ROI activity in infants and children. Indeed, skull conductivity varies substantially across individuals and age groups (McCann et al., 2019). The EEG templates could be refined by adjusting this parameter to best match the data.

## Acknowledgements

This work was supported by BBSRC grant BB/N018516/1 and Wellcome Trust ISSF 204821/Z/16/Z.

