## Supplementary material for "Estimating functional EEG sources using topographical templates"

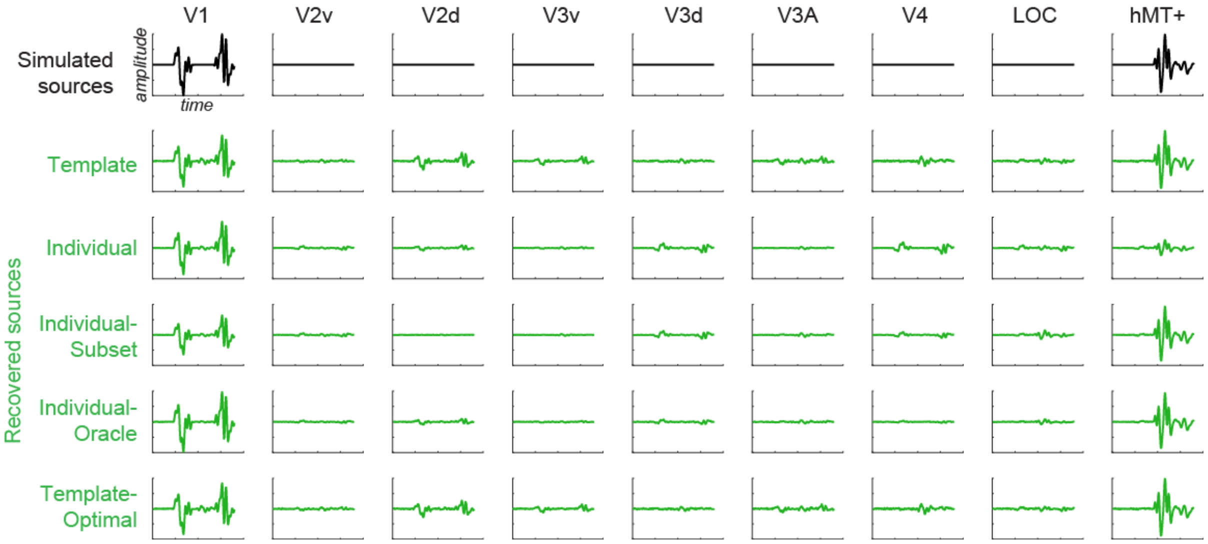


*Figure S1. Simulation of V1 and hMT+ activity over time (in black) and the retrieved signal (in green) from EEG scalp responses using various source localization methods.*

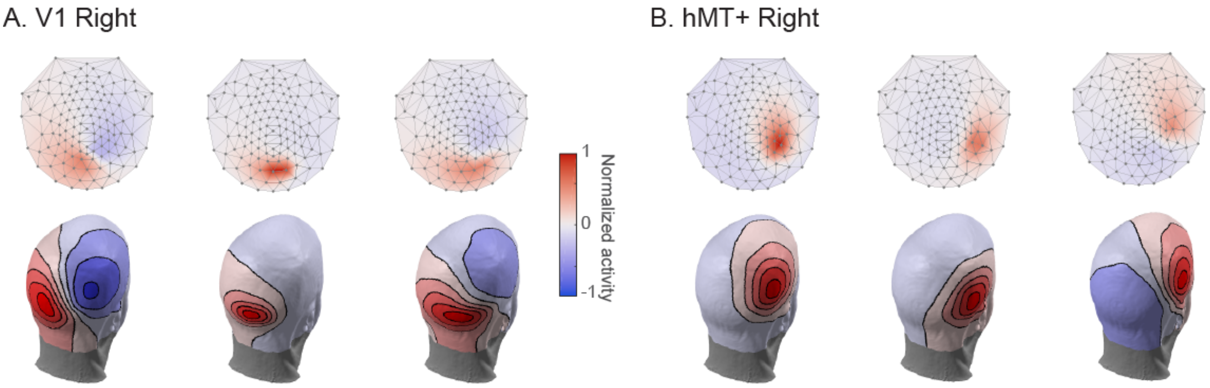


*Figure S2. Illustration of individual differences in EEG scalp responses. 2D (top) and 3D (bottom) representation of the scalp activity for V1 (A) and hMT+ (B) in the right hemisphere for three different individuals.*

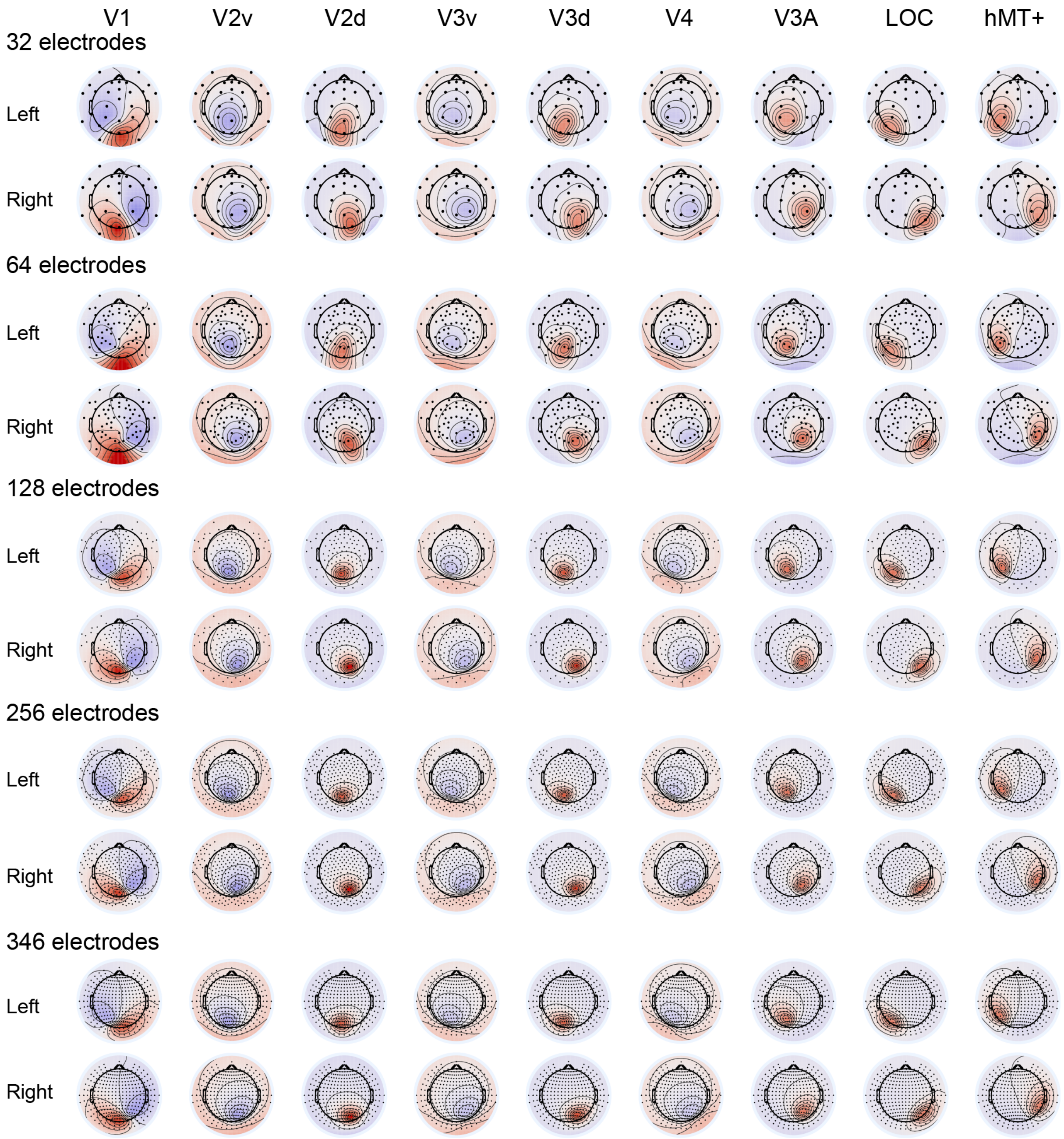


*Figure S3. EEG templates for a standard EGI-system with 32, 64, 128, 256 electrodes and for a standard 10-05 system with 346 electrodes. The intensity of the color indicates the amplitude of positive (red) and negative (blue) activity.*

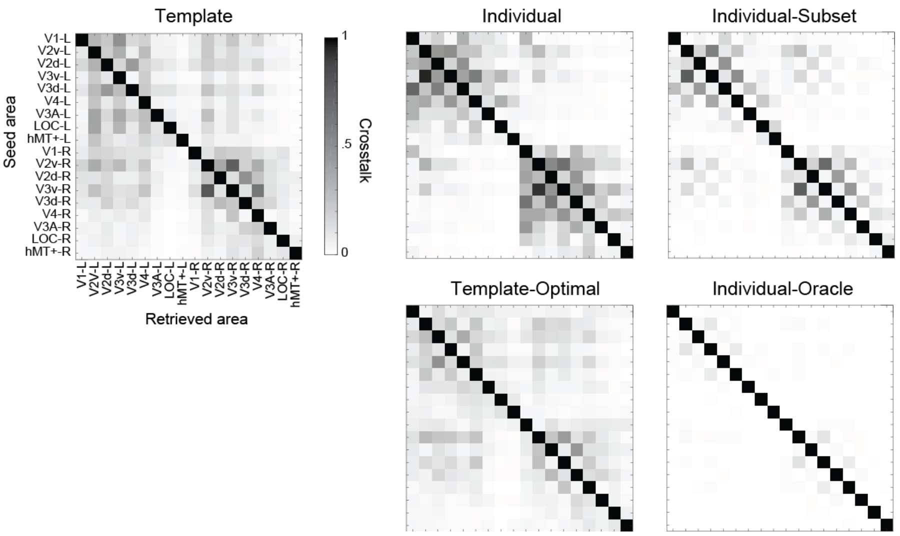


*Figure S4. Crosstalk (leakage) between ROIs for different source localization methods. The amount of crosstalk (normalized for each ROI; per row) was calculated for an EEG signal simulated using a 128 EGI montage with an SNR of 200 and averaged across 50 individuals. The darker the square, the more crosstalk between those two areas.*

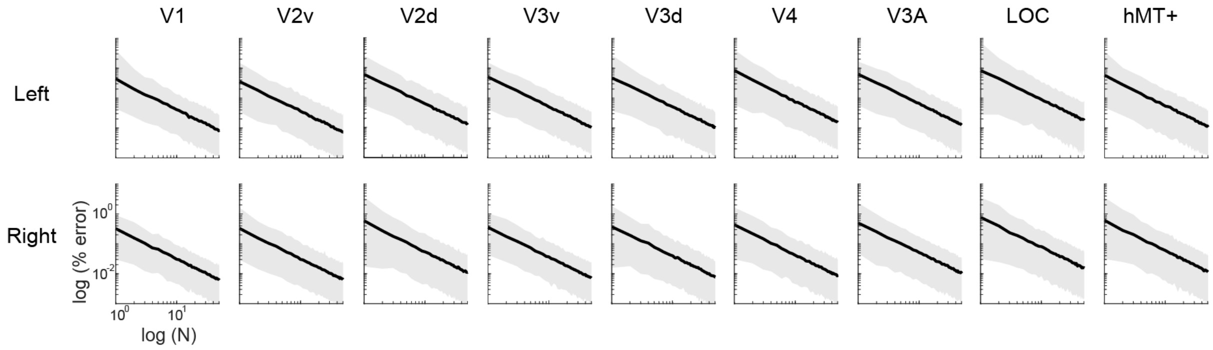


*Figure S5. The variability in EEG templates (% error) is inversely proportional to the number of participants. In log-log plots, this exponential decay function follows a 1/N slope with the intercept determined by the mean sample at N=1. The shaded area represents 95% confidence interval.*

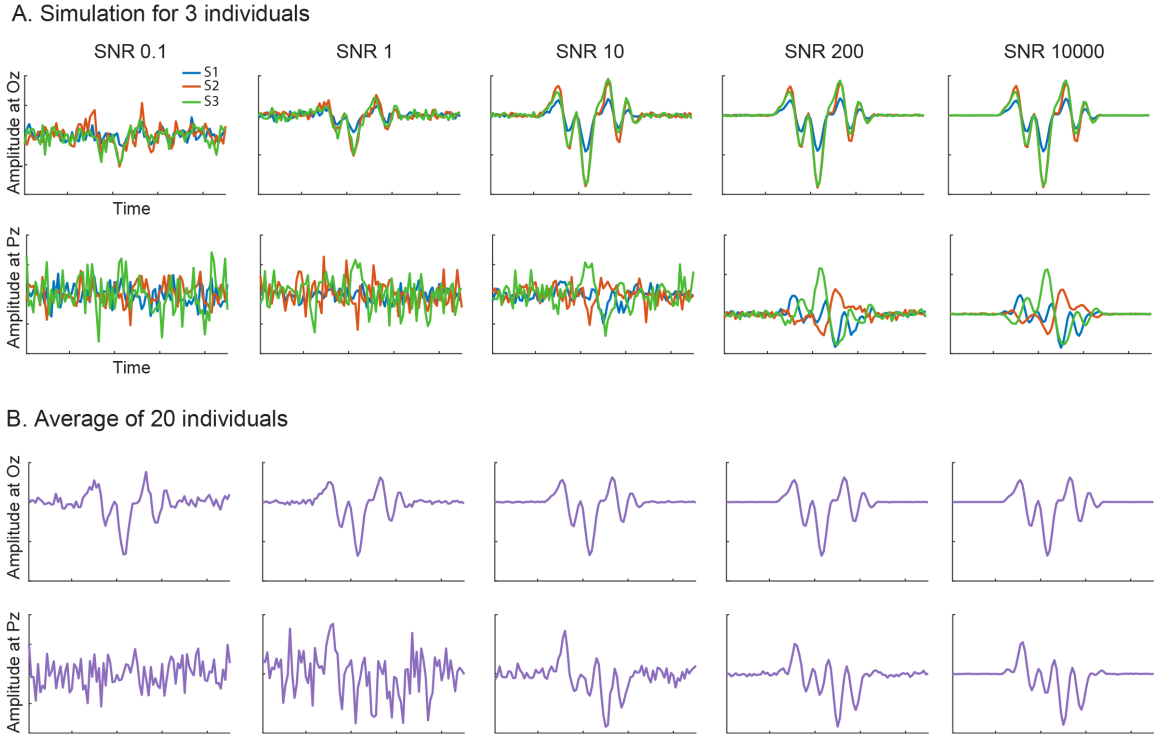


*Figure S6. Simulation of ERPs with different levels of SNR for three individuals (A) and the average ERP of 20 individuals (B) at two electrodes location (Oz and Pz). Note the variation across participants and electrodes in A. Typical recorded data will be in the 10-200 SNR range when averaged across participants (B).*

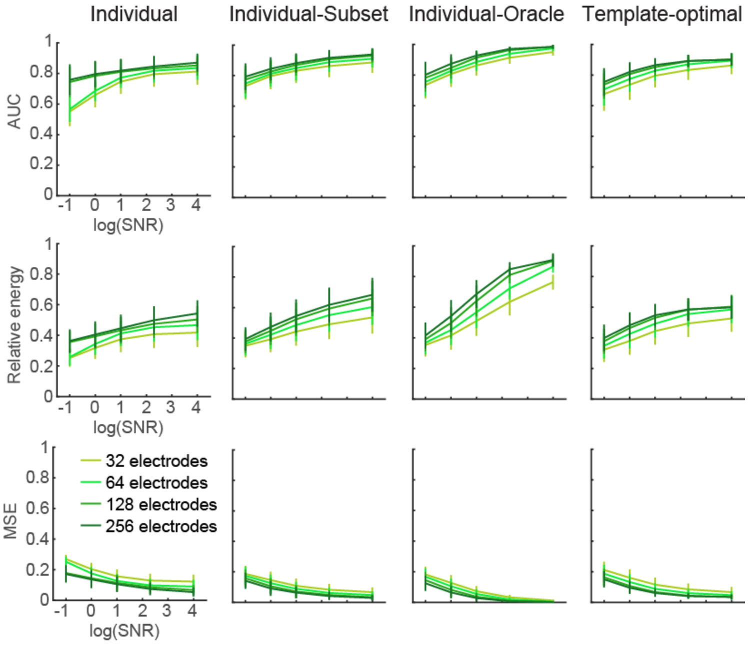


*Figure S7. Source localization performance using different methods and different EEG montages with 32, 64, 128 and 256 electrodes. Brain sources are recovered from the simulation of two bilateral ROIs chosen randomly.*

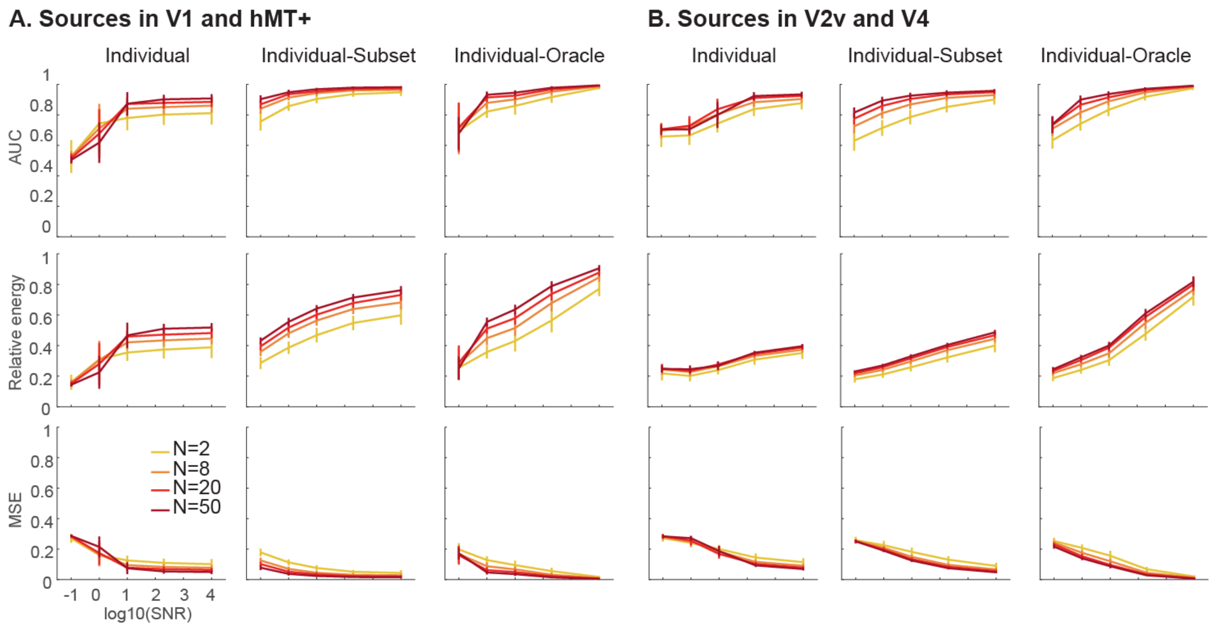


*Figure S8. Source localization performance with a 128 electrodes EGI montage at different SNR levels for N=2, 8, 20 or 50 participants using different source localization methods with L-curve regularization. Sources are simulated bilaterally in V1 and hMT+ (A) or V2v and V4 (B).*
